# Detection of protein symmetry and structural rearrangements using secondary structure elements

**DOI:** 10.1101/2025.09.18.677065

**Authors:** Runfeng Lin, Sebastian E. Ahnert

## Abstract

Many proteins exhibit a degree of internal symmetry in their tertiary structure, including circular permutations. These characteristics play an important role in terms of the functional robustness of proteins against mutations, and are pivotal for the study of protein function and evolution. Proteins exhibiting internal symmetry often demonstrate enhanced functional benefits, such as increased binding affinity due to repeated structural motifs, and evolutionary advantages that may result from gene duplication or fusion events, leading to more robust and adaptable molecular architectures. Similarly, circular permutations have been linked to improved catalytic activity and enhanced thermostability, opening new avenues in the design of enzymes and other functional proteins. In this study, we introduce a novel computational pipeline that leverages secondary structure elements (SSEs) as a compressed yet informative representation of protein structure to detect both symmetry and structural rearrangements effectively. Our method shows strong advantages against the Combinatorial Extension with Circular Permutations (CECP) for computational cost and identifies 17,130 circularly related proteins. Additionally, it detects 26,739 proteins associated with indel mutations. Notably, 8,855 of these exhibit both circular permutation and indel mutations-highlighting a potential co-evolutionary relationship between these two types of structural variation. We also recovered most symmetric proteins identified by the CE-symm tool and revealed more than 2.5 times as many symmetric proteins in the same dataset. By integrating precise boundary detection, clustering analysis, and rigorous cross-validation against established tools, our framework robustly maps internal structural features. These insights deepen our understanding of protein architecture and evolution, and can also inform de novo protein design and engineering.

## 1 Introduction

Proteins fulfil a myriad of essential functions in every biological cell, and their three-dimensional (3D) structures underpin these functional roles. The tertiary structure of a protein is determined by the intricate interplay between amino acid interactions and environmental influences. While significant progress has been made in predicting protein structures [1] and simulating protein dynamics [2], many structural studies have primarily focused on global folding patterns, often overlooking finer internal characteristics such as symmetry, indel mutations, and circular permutations [3, 4].

Research has demonstrated that symmetric and simplified structural motifs are intrinsically more likely to emerge in evolutionary processes [5]. Internal symmetry in proteins is more than a mere structural curiosity; it has profound functional implications. Proteins with symmetrical architectures can exhibit increased binding affinities due to the presence of multiple identical binding sites, and they may also benefit from enhanced stability and robustness as a result of evolutionary events like gene duplication or fusion [6]. Circular permutations, wherein the protein chain is rearranged so that the original termini are reconnected at new positions, have been associated with increased catalytic efficiency and improved thermostability [7, 8]. Insertions and deletions (indels) are a crucial force in protein evolution. By subtly altering the length and flexibility of loop regions, indels can modify protein function—affecting binding interactions, enzyme specificity, and overall stability. Although typically small, these changes offer a mechanism for evolutionary innovation, allowing proteins to adapt new functions while preserving their essential structural framework [9]. Such rearrangements suggest that nature exploits these features to fine-tune protein function and adaptability.

Recent advancements have shown that secondary structure elements (SSEs) provide a powerful, compressed representation of protein structures. (Example in Figure 1. More details are available in Method 4.1) By focusing on SSEs, one can reduce the complexity inherent in residue- or atomic-level representations of protein structure while retaining critical information about the protein’s overall architecture [10].

**Figure 1:**
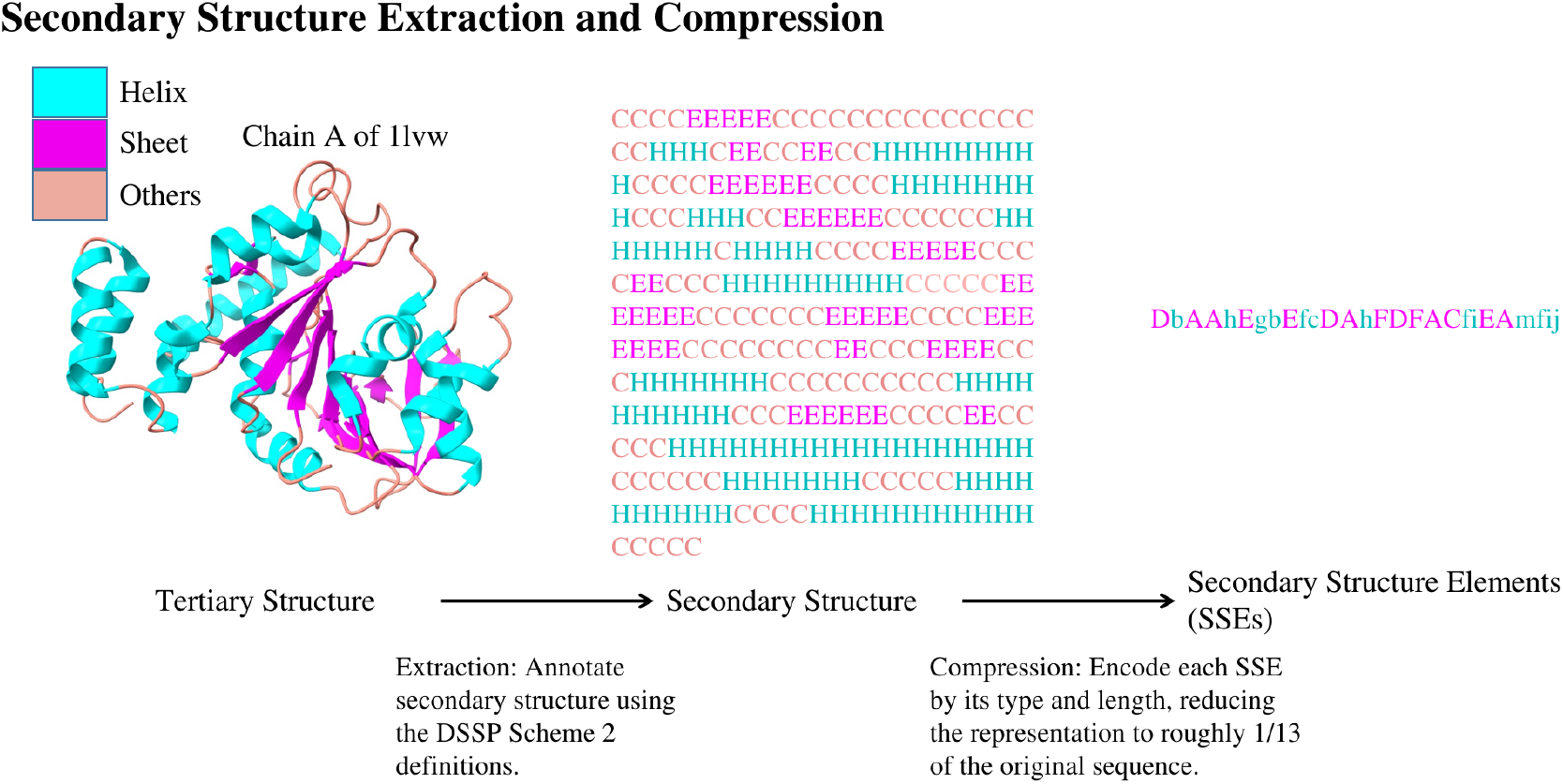
Secondary structure elements extraction. Secondary structures are first extracted from tertiary structure using DSSP [11] based on scheme 2, and then further compressed into Secondary Structure Elements (SSEs) [10].

In this work, we used SSEs to develop a pipeline that detects protein symmetry, circular permutations, and indel mutations. Our approach begins with the extraction of SSEs, followed by precise boundary detection and clustering algorithm designed to capture subtle rearrangements. The pipeline is validated against established databases and benchmarked against state-of-the-art tools, ensuring that our method is both reliable and sensitive to variations in protein structure. In terms of speed, our SSE-based method is significantly outperforms the Combinatorial Extension with Circular Permutations (CECP) [4], which is more than 16,000 times slower. Our approach uncovers 17,130 circularly related proteins. Notably, over 8,855 of these are both circularly permuted and exhibit evidence of indel mutations. This suggests that circular permutations may often occur in tandem with indel mutations. We also recover most of the symmetric proteins that can be identified using the CE-symm tool [12] and reveal in total more than 2.5 times as many symmetric proteins in the same dataset, underscoring the sensitivity and breadth of the pipeline.

## 2 Results

### 2.1 SSE Duplication Pipeline

In order to detect internal symmetries we developed an SSE-based pipeline to search for structural duplications. We evaluated this SSE-duplication pipeline on a non-redundant subset of the RCSB PDB, comprising 71,781 protein chains (sequence identity ≤ 90%, length ≥ 5 SSEs). This filtering minimizes bias and ensures broad structural coverage. Figure 2(a) summarizes the three main steps: SSE duplication, precise boundary detection, and validation.

**Figure 2:**
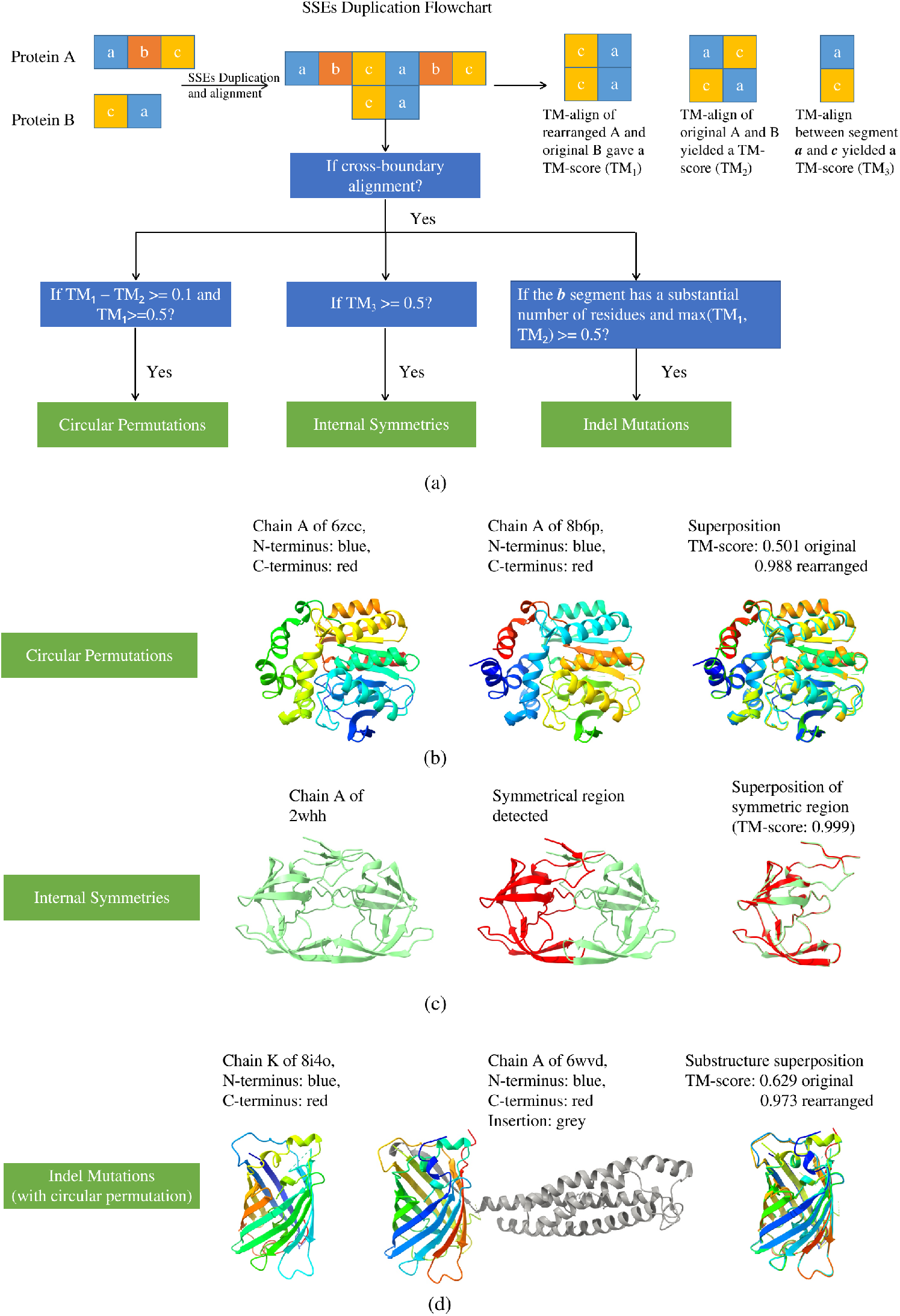
SSE Duplication Workflow and Applications. (a) Workflow of the SSE duplication pipeline. (b) Application to circular permutation detection. (c) Application to circular permutation with indel mutation detection. (d) Application to symmetric region detection. Our SSE duplication pipeline demonstrates broad applicability and robust performance across multiple types of protein structural property detection.

#### 2.1.1 Detection of Circular Permutations

Applying the pipeline to this dataset, we identified 67,984 circularly permuted pairs across 17,130 unique proteins (Figure 2(b)). Leiden clustering [13] grouped these proteins into 453 communities, with 70% of members concentrated in the ten largest clusters—indicating a highly skewed distribution.

For external validation, we assessed the performance of SSE duplication against the Combinatorial Extension with Circular Permutations (CECP) [4] method using the same dataset. As CECP is computationally intensive—requiring an estimated 2.2 million CPU hours for full-scale analysis—we restricted the comparison to the 67,984 pairs detected by SSE duplication. In addition, we used the Circular Permutation DataBase (CPDB) as a representative benchmark set and compare the how well CECP and SSEs duplication can reproduce the results.

*Comparison with CECP* – Among the 67,984 pairs analyzed using CECP, only 40,058 had a TM-score above 0.5, and not all were recognized as circular permutations. In contrast, SSE duplication requires both a post-rearrangement TM-score above 0.5 and an improvement of at least 0.1 over the original to qualify as a circular permutation. CECP identified 53,690 pairs that contain cross-boundary alignment in total, but only 22,758 of these met the TM-score threshold after rearrangement. Notably, the SSE-duplication pipeline completed the entire dataset in approximately 130 CPU hours, making it over 16,000 times more efficient than CECP. For a detailed comparison see Table 1.

**Table 1:** Comparison of SSE-duplication and CECP on a filtered RCSB dataset.

| Metric | SSE Duplication | CECP |
| --- | --- | --- |
| *Overall dataset* |  |  |
| Unique proteins analysed | 17,130 | 17,130 |
| Cross-boundary alignment (pairs) | 67,984 | 53,690 |
| Pairs with TM-score $\geq 0.5$ | 67,984 | 40,058 |
| Pairs with TM-score $\geq 0.5$ and $TM_{\text{SSE}} - TM_{\text{CECP}} \geq 0.1$ | 67,984 | 22,758 |
| *Pairs identified by both methods* |  |  |
| Highly similar (TM-score $\geq 0.8$) | 4,433 | 4,663 |
| Pairs where $TM_{\text{SSE}} - TM_{\text{CECP}} > 0.1$ | 7,600 | — |
| Pairs where $TM_{\text{CECP}} - TM_{\text{SSE}} > 0.1$ | — | 1,018 |
| *CPs unique to SSE duplication* |  |  |
| Unique SSE-only CP pairs | 14,294 | — |
| Highly similar (TM-score $\geq 0.8$) | 236 | 217 |
| Pairs where $TM_{\text{SSE}} - TM_{\text{CECP}} > 0.1$ | 4,095 | — |
| Pairs where $TM_{\text{CECP}} - TM_{\text{SSE}} > 0.1$ | — | 660 |

We further analyzed the results by dividing them into two groups: (a) circular permutations (CPs) identified by both methods, and (b) CPs uniquely identified by the SSE duplication method.

For the subset identified by both methods, the SSE duplication pipeline generally recovers substructures with higher overall similarity. The number of highly similar substructures is comparable between the methods (TM-score ≥ 0.8): 4, 433 for SSE duplication versus 4, 663 for CECP. For the same protein pairs, SSE duplication identifies over 7, 600 pairs with markedly higher similarity than CECP, defined by

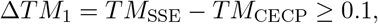

whereas CECP exceeds SSE in only 1, 018 pairs,

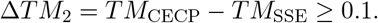

For CPs not recognized by CECP, we observed the same trend. SSE duplication continues to identify more highly similar substructures, with 236 cases exceeding a *TM* -*score* of 0.8, compared to 217 for CECP. Again, in terms of relative similarity, SSE identifies 4,095 pairs with Δ*TM*_1_ *>*= 0.1, while CECP identifies only 660 such cases with Δ*TM*_2_ *>*= 0.1 .

Upon visual inspection of several CECP pairs that extracted structurally similar substructures but were not classified as CPs, we found that many involve indel mutations combined with near-symmetric rearrangements. In particular, we observed cases where a smaller protein aligns well with both a continuous internal region and the disjoint N- and C-termini of a larger protein. More details of these comparisons can be viewed at Table 1 and an example of this phenomenon is shown in Figure S2.

*Comparison with CPDB* – For the benchmark comparison, we applied both the SSE-duplication method and CECP to reproduce circular-permutation pairs reported in the Circular Permutation

DataBase (CPDB). To ensure a fair comparison for the SSE-based approach, we excluded any pair in which either protein contains fewer than five secondary-structure elements (SSEs), yielding a filtered set of 3,102 pairs. Within this set, the SSE-duplication method identified 2,410 pairs with cross-boundary alignments, whereas CECP identified 2,853 pairs.

Among the pairs identified by both methods, CECP generally reported higher structural similarity between the extracted substructures; however, the effect size was modest, with an average TM-score increase of ≈ 0.047 relative to the SSE-duplication method, i.e.,

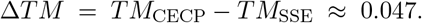

See Fig. 3.

**Figure 3:**
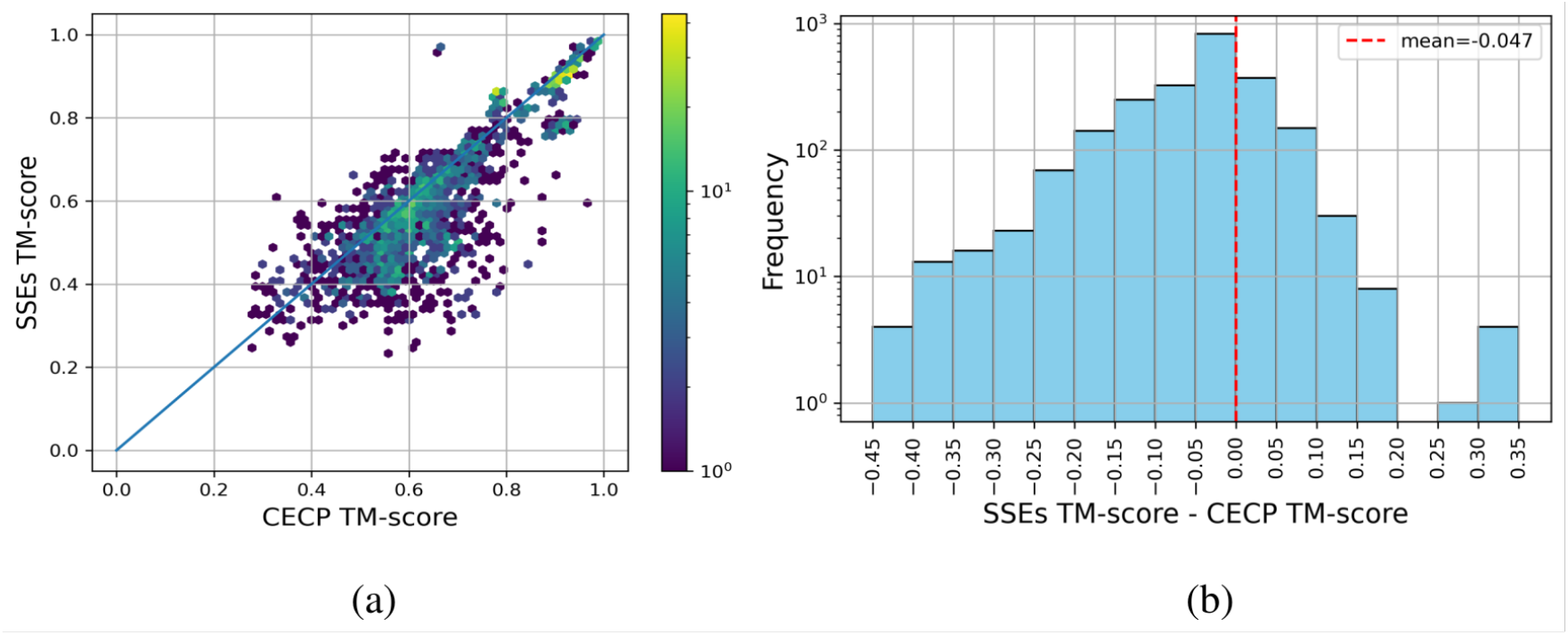
Comparison of SSE Duplication and CECP Performance in CPDB. (a) Correlation between TM-scores after structural rearrangement using SSE-based duplication and CECP methods. (b) Histogram of the difference between post-rearrangement TM-scores from SSE duplication and CECP (SSE − CECP). Overall, the two methods yield comparable structural similarity scores, with CECP performing slightly better on average than SSE duplication.

Given the low computational cost of the SSE-duplication method, we performed exhaustive pairwise comparisons among proteins in the CPDB dataset. This analysis uncovered an additional 1,559 protein pairs that satisfy the criteria for circular permutation but were not previously annotated in CPDB. These newly identified pairs are efficiently detected by the SSE-duplication method. In contrast, identifying these pairs with CECP would require substantially more computational resources, underscoring the superior scalability and practical sensitivity of our approach.

#### 2.1.2 Indel Mutations

As many novel circularly permuted pairs exhibited substantial gaps between their aligned boundaries, we extended our analysis to indel mutations. We found that there are in total 199,804 pairs of proteins that are indel mutants of each other, corresponding to 26,379 unique proteins. Furthermore 15,966 of the previously identified 67,984 permuted pairs, corresponding to 8,855 unique proteins, also harbour indels. These dually modified proteins cluster into 813 groups, highlighting the additional structural diversity introduced by insertions and deletions (Figure 2(c)). Notably, many fluorescent proteins were identified to contain both circular permutations and indel mutations which is consistent with the known tolerance of green flourescent protein (GFP) to both circular permutations and indel mutations [14].

#### 2.1.3 Symmetry detection from cross-boundary alignment

Many hits from the SSE-duplication pipeline exhibited a TM-score difference (ΔTM) below 0.1 between the original and permuted orders, and thus were not classified as circular permutants. This small difference suggests that such proteins may possess inherent structural symmetry or exhibit domain-size imbalance that biases alignments toward the same region. Visual inspection supports this interpretation, revealing frequent symmetric features. To systematically assess symmetry, we split each protein at its duplication boundary and calculated the TM-score between the resulting segments; proteins with *TM* ≥ 0.5 were classified as symmetric [15]. For comparison, we also applied CE-Symm [12] to the same non-redundant dataset, selecting proteins with *TM* ≥ 0.5 and more than one repeat unit. Although the number of symmetric proteins detected by SSE-duplication (7,226) and CE-Symm (7,108) was similar, the overlap between the two methods was limited, with a Jaccard index of just 0.240. The unique predictions made by each method are illustrated in Supplementary Figure S1.

Comparing unique predictions highlights complementary strengths:

- **CE-Symm** excels at detecting proteins whose repeat units have very high structural similarity.
- **SSE-duplication** uncovers symmetry in proteins with more complex SSE arrangements (See supplementary Figure S3).

In general, the unique predictions by these two methods have very similar distribution of symmetry and number of SSEs. The SSE-duplication method is currently limited by its reference set for cross-boundary alignments and support for only two-fold symmetry.

Together, these results demonstrate that our SSE-duplication pipeline not only reliably captures standard circular permutations with strong agreement to CPDB and provides a comprehensive view of structural symmetry in proteins, but also uncovers novel, indel-modified permuted proteins—highlighting its sensitivity and broad applicability to protein structural variation.

### 2.2 Self-Scanning of SSEs

To overcome the limitations of detecting symmetry solely based on cross-boundary alignment, we employed a scanner-based self-alignment of SSEs.(see Figure 4 more details can be found at Method 4.5)

**Figure 4:**
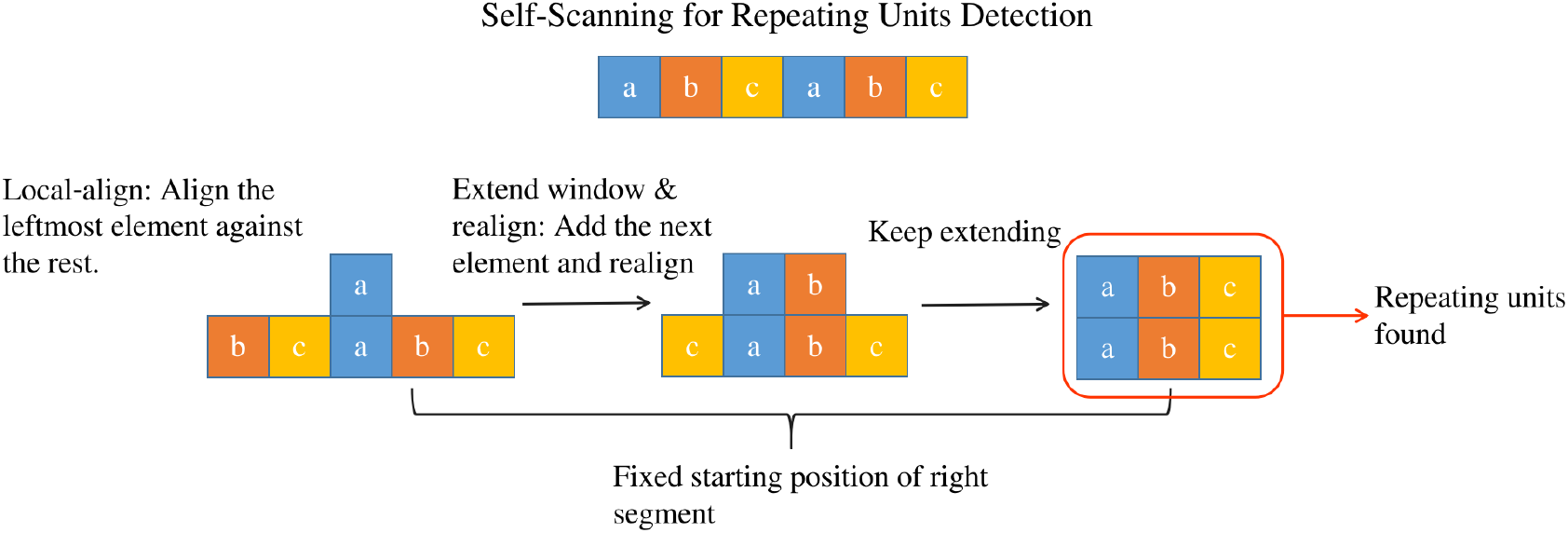
Self-Scanning of SSEs. Starting from the leftmost secondary structure element (SSE) in each sequence, we slide a window that initially contains three SSEs and extend it rightward one SSE at a time until only three SSEs remain at the end. Each extracted segment pair is then verified using TM-align to assess internal symmetry.

**Figure 5:**
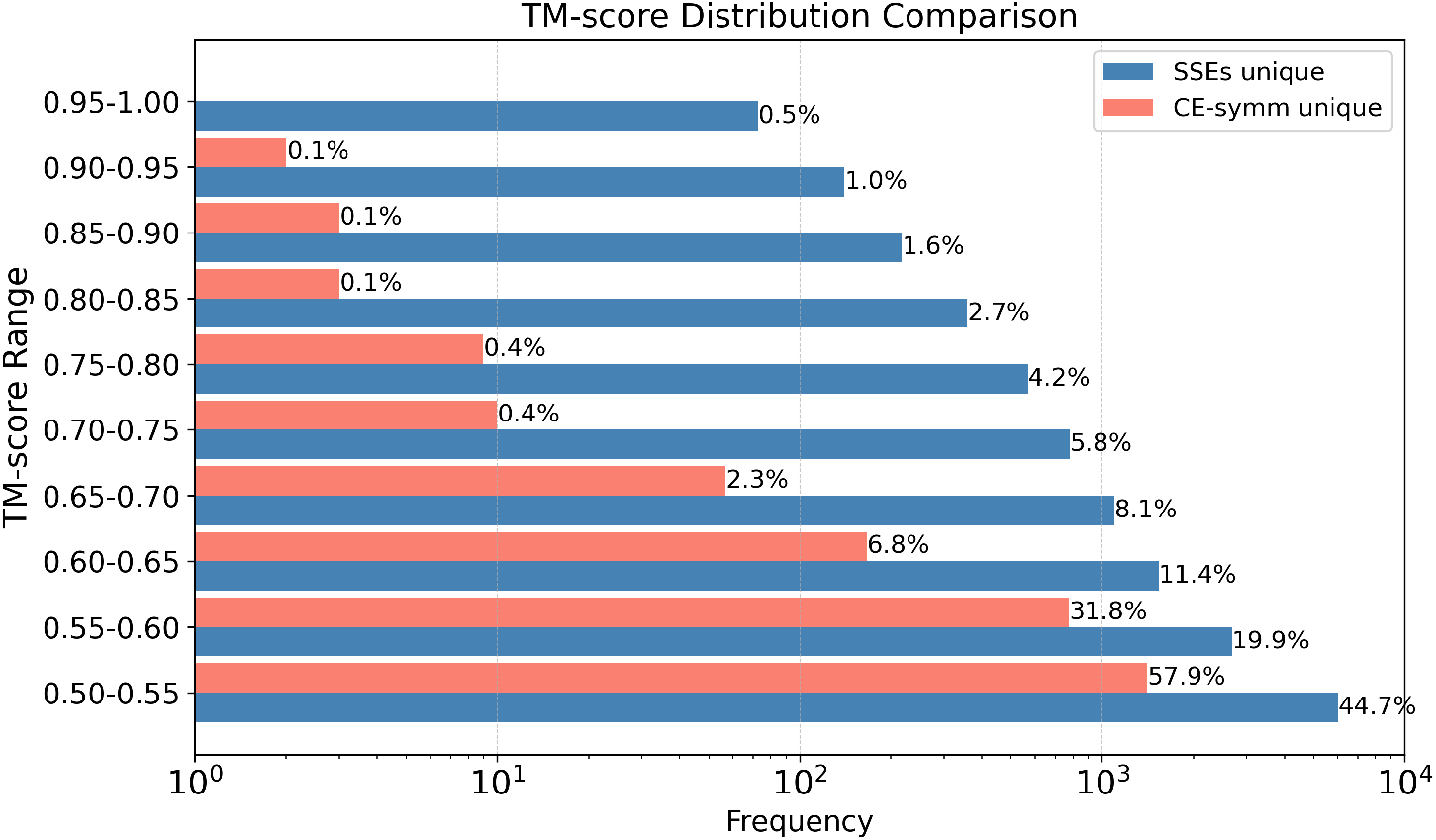
TM-Score Distribution of Unique Predictions. This distribution compares the TM-scores of unique symmetry protein predictions by the SSE-based and CE-Symm methods. Nearly 90% of the CE-Symm predictions lie between 0.5 and 0.6, suggesting moderate symmetry, while the SSE-based method identifies a broader range, capturing many highly symmetric structures.

This approach identified 18,173 proteins that contain sub-structure pairs with TM-scores above 0.5 (requiring a minimum of 3 SSEs per substructure) and detected more symmetric proteins than the union of those identified by the CE-Symm and SSE-based methods for circular permutations.

Specifically, this method produced 13,505 unique findings and recovered 65.6% of the symmetric proteins found by CE-Symm, which had 2440 unique predictions. Comparison of the TM-scores in the symmetric regions of the unique predictions reveals that nearly 90% of the CE-Symm unique predictions have TM-scores below 0.6, whereas only 65% of the SSE-based unique predictions fall below that threshold. This indicates that CE-Symm may miss many highly similar, symmetric structures, while the SSE-based method tends to omit structures with only moderate similarity.5

Furthermore, analysis of the correspondence between the number of SSEs in a protein and the TM-score of its symmetric region shows that CE-Symm struggles to capture local symmetry in large proteins—none of its unique predictions contains more than 50 SSEs—whereas the SSE-based method can detect symmetric regions in proteins with up to 500 SSEs. Many of these large proteins exhibit extremely symmetric structures. These findings highlight the superior sensitivity and scalability of the SSE-based approach in detecting local structural symmetry. It not only recovers more symmetric proteins overall but also captures highly symmetric regions in large and complex structures that CE-Symm consistently misses. (See Figure 6)

**Figure 6:**
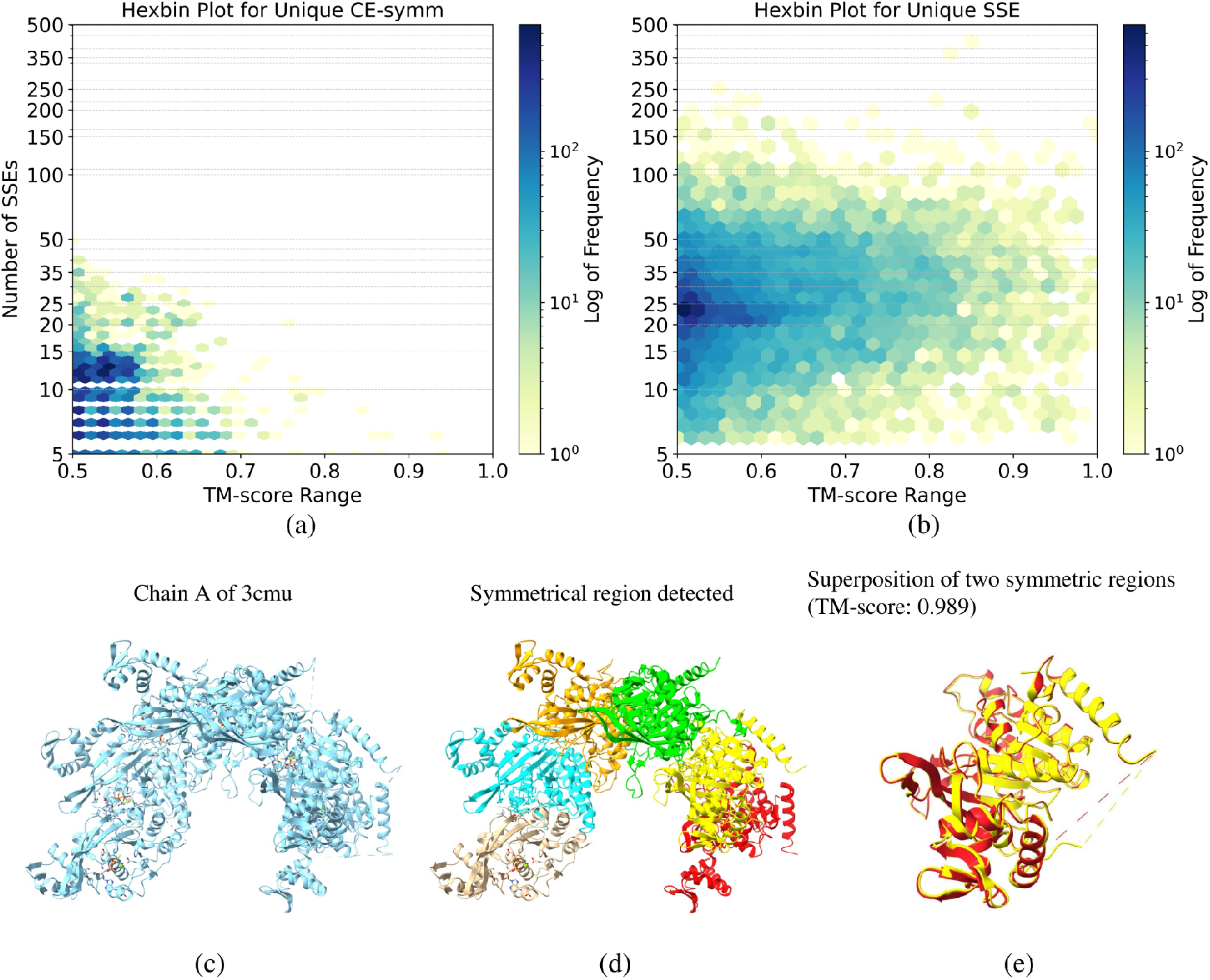
TM-Score and Number of SSEs Correspondence of Unique Predictions. (a) TM-score vs. number of SSEs for proteins uniquely detected by the SSE-based method. (b) Same for CE-Symm. (c) A representative example of a large and highly symmetric protein identified by the SSE-based method but missed by CE-Symm. These plots show that CE-Symm fails to detect symmetric regions in large proteins (none with above 50 SSEs), whereas the SSE-based approach successfully identifies local symmetry even in proteins with up to 500 SSEs, demonstrating superior sensitivity and scalability for detecting structural symmetry.

**Figure 7:**
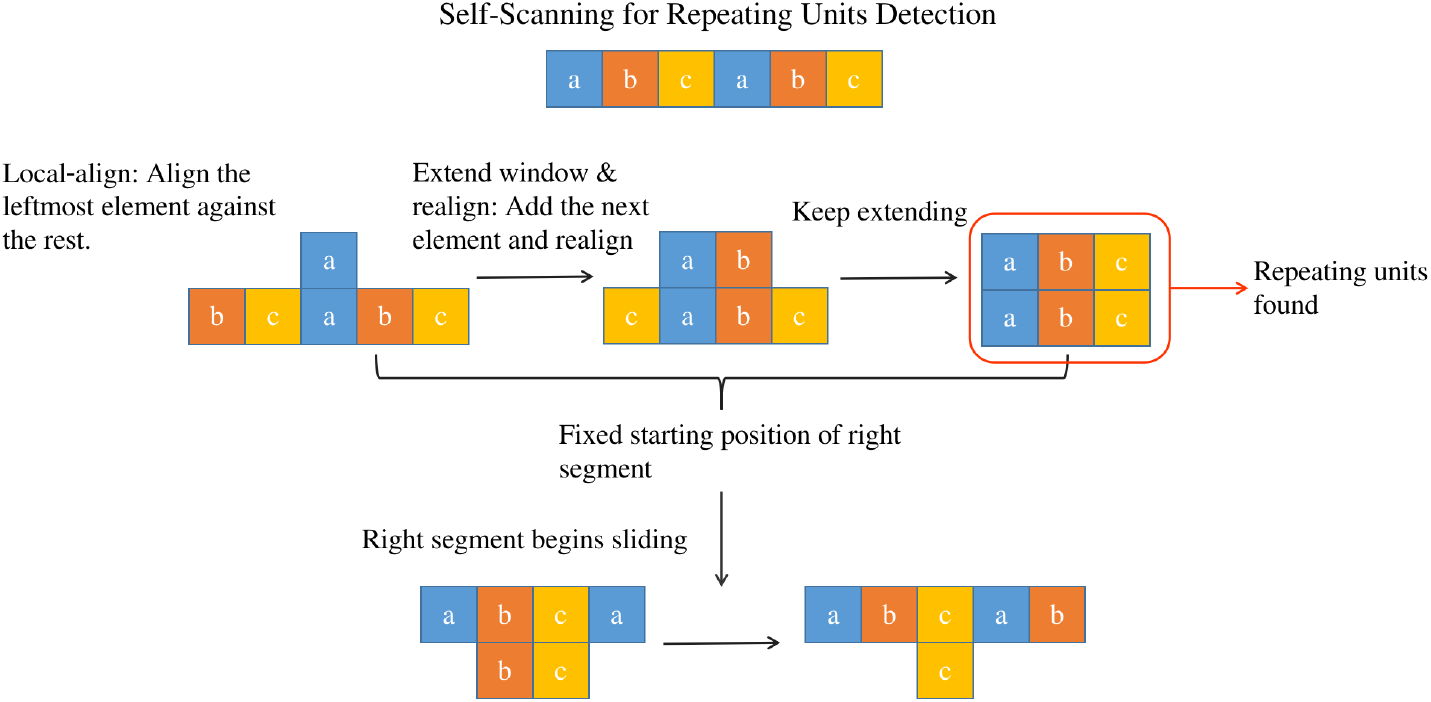
Self-Scanning for Repeating Units Detection. Illustration of the self-scanning method for detecting repeating structural units. The input sequence is scanned by fixing the starting position of the left segment, extending it one element at a time, and aligning it against the right segment. When an optimal alignment is found (highlighted), repeating units are identified. After all fixed positions of the right segment are exhausted, the right segment begins to shift, continuing the scan.

We further compared the number of repeating units detected by each method across the overlapping set of proteins. In most cases, both methods identified only two symmetric units, and their results were generally consistent. However, significant discrepancies emerged when proteins contained a larger number of repeats. In such cases, CE-Symm typically identified more repeated units than the SSE-based method. (See Supplementary Figure S5)

Visual inspection of these differences revealed that CE-Symm is more effective at detecting smaller, localized repeating units, particularly in highly symmetric regions. This discrepancy arises from the SSE-based method’s summarization approach, which may overlook smaller units. For example, consider a protein composed of four identical repeats (A-A-A-A). During self-scanning, the first repeat may align better with the third than with the second. Because the scanning window extends one SSE at a time, the boundaries between units may shift such that the four repeats are grouped into two larger units, rather than being recognized individually. More details on the self-scanning procedure and how repeated units are summarized are provided in the Methods section 4.5.

## 3 Discussion and Conclusion

In this study, we developed an SSE-duplication pipeline to effectively detect internal structural features in proteins by aligning across secondary structure element (SSE) boundaries. The pipeline successfully recovered most circular permutations previously annotated in redundant RCSB entries and the CPDB, additionally identifying 1,559 previously unannotated pairs within CPDB. In total, we detected 67,984 circularly permuted pairs across 17,130 unique proteins out of 71,781 analyzed. A recognized limitation is that proteins with few SSEs or a TM-score below 0.6 might evade detection due to weakened alignment signals from sparse SSE representations.

Significantly, our analysis revealed a notable intersection between circular permutations and insertion-deletion (indel) mutations. Among the analyzed proteins, 26,379 proteins exhibited indel mutations, comprising 199,804 unique indel-mutant pairs. Strikingly, 8,855 proteins exhibited both circular permutations and indel mutations, emphasizing a potential co-evolutionary relationship between these two structural variations. Network analysis further revealed highly skewed cluster-size distributions in both the full circular permutation dataset and the subset containing indel mutations, reinforcing the hypothesis of co-evolutionary mechanisms linking circular permutation and indel events.

To assess internal symmetry, we compared the results of cross-boundary SSE alignments against the established CE-Symm method. Although both methods yielded similar counts (7,226 proteins for SSE duplication versus 7,108 for CE-Symm), the overlap was limited (Jaccard index = 0.24), indicating complementary strengths. CE-Symm excels at identifying small, highly similar repeat units, whereas our SSE-duplication method covers a broader range of SSE counts but remains inherently restricted to detecting twofold symmetry and relies on boundary crossings or database matches.

Despite the success of our SSE-duplication approach, it has inherent constraints. The method requires sufficiently long SSEs and cannot resolve repeats involving more than two units. Additionally, the reduction of full protein structures to SSEs may obscure flexible loops and small helices, thus reducing sensitivity in regions with sparse or short SSEs. Future enhancements could involve incorporating additional structural features, such as backbone geometry or loop conformations, into the SSE representation. An adaptive substitution matrix implemented via a neural network could further improve circular permutation boundary detection, though both strategies would increase computational complexity.

To address limitations inherent to SSE duplication for internal symmetry detection, we developed a self-scanning SSE approach specifically tailored for this purpose. This method significantly outperformed CE-Symm, identifying 18,173 symmetric proteins and covering 65.6% of CE-Symm’s predictions. Missed cases were primarily associated with proteins containing few SSEs or moderate structural similarity..

Further analysis indicated that the SSE-based method surpasses CE-Symm not only in the number of detected symmetric proteins but also in handling structurally complex proteins. Our approach successfully identified symmetry in proteins containing up to 500 SSEs, whereas CE-Symm’s capability is restricted to fewer than 50 SSEs.

Regarding repeated unit identification, both methods generally agreed, though CE-Symm demonstrated greater proficiency in capturing spatially independent units. This discrepancy arises from differing methodologies in defining and summarizing repeats, as detailed in the Methods section.

Additionally, CE-Symm uniquely distinguishes between open symmetry (linear or helical repeats) and closed symmetry (cyclic or looped arrangements), a feature not supported by our SSE-based approach. Nevertheless, the SSE-duplication pipeline offers significant advantages in computational efficiency, scalability, and sensitivity, providing robust opportunities for advanced protein structural analysis and protein engineering applications.

## 4 Method

The alignment and substitution matrix for SSEs is based on the SSE-based structure searching pipeline [10]. The filtered RCSB dataset was generated by applying a 90% sequence identity threshold to minimize redundancy.

### 4.1 Secondary structure elements (SSEs) and alignment

The secondary structure of each protein was first assigned using DSSP [11] (Scheme 2), categorizing residues into three classes: Random Coil (C), Beta Strand (E), and Alpha Helix (H). Consecutive residues of the same class are then grouped into secondary structure elements (SSEs), labeled by type and length—for example, HH-a, HHH-b, or EE-A (Figure 1). In terms of Shannon entropy, the SSE representation is 9.85-fold more compact than the original amino acid sequence.

Pairwise alignment of SSEs is performed with the Smith–Waterman algorithm using element-specific gap penalties, and the substitution matrix was optimized via a genetic algorithm to maximize recovery of SCOPe domains [16]. The resulting alignment features are fed into a hybrid ResNet–BiLSTM model to predict TM-scores. For a full description of the SSE-based structure-searching pipeline, see Lin et al. [10]

### 4.2 Duplication of SSEs

Circular permutation and indel mutations are determined by cross-boundary alignments, achieved by duplicating the query SSEs and aligning them to the original target SSEs. Illustration of these alignments can be found at Figure 2 (a). Alignments crossing the original protein boundary are filtered by a predicted TM-score threshold of 0.5 and further verified by actual TM-score calculations. Circular permutations with indel mutations are defined as cases where a gap exists between cross-boundary segments. This gap is typically defined as at least 40 residues (approximately three SSEs, based on the requirement of at least three SSEs per side) or at least 60% of the length of the smaller segment in terms of residues. Indels of 20–30 residues are generally treated as minor loop variations [17]. Symmetry is determined if cross-boundary aligned segments achieve a TM-score above 0.5 [15].

### 4.3 TM-align

TM-align [18] is a pairwise protein-structure alignment algorithm that uses only the C_*α*_ coordinates of two protein chains to compute a normalized TM-score [19] between 0 and 1. A TM-score above 0.5 generally indicates that two structures share the same fold [15].

For circular permutation analysis, two proteins with identical C_*α*_ coordinates and the same number of residues in each permuted segment will yield a TM-score of approximately 0.5 in the original sequence order, rising to 1.0 after correctly rearranging the permuted units. In practice, the pre- and post-rearrangement TM-scores depend not only on the overall structural similarity but also on the relative size of the largest permuted segment.

### 4.4 Symmetry Proteins by CE-Symm

CE-Symm is a dedicated tool for detecting internal symmetry in proteins [12]. It applies the Combinatorial Extension (CE) algorithm to align a protein with a copy of itself while forbidding near-diagonal alignments to avoid trivial matches. In this study, we consider symmetric segments with TM-scores above 0.5 with more than 1 repeated units as symmetric proteins.

### 4.5 Self-Scanning of SSEs for Symmetry Detection

To detect symmetric proteins, we implemented a self-scanning SSE alignment strategy. Starting from the first SSE, a window of three consecutive SSEs is aligned against the remainder of the chain; the window then extends one SSE at a time to the right while its start point stays fixed, stopping when fewer than three SSEs remain. Since RCSB proteins contain on average ∼ 19 SSEs, this entails 14 alignments per protein, ensuring scalability to large datasets. Because self-scanning generates many alignments, we filter for those with TM-score *>* 0.5, and identify the minimal repeating-unit length by recording how far the right-segment start shifts between successive optimal alignments (see Figure 1). To consolidate redundant hits—alignments that share similar or fully overlapping boundaries—we construct a graph in which nodes are segments and edges join segments with high structural similarity. We then extract connected components and compute each component’s repeat count as the total boundary coverage divided by its minimal unit length.

Further implementation details are given in Algorithm 1.

#### Algorithm 1

Self-Scanning SSE Repeat Detection

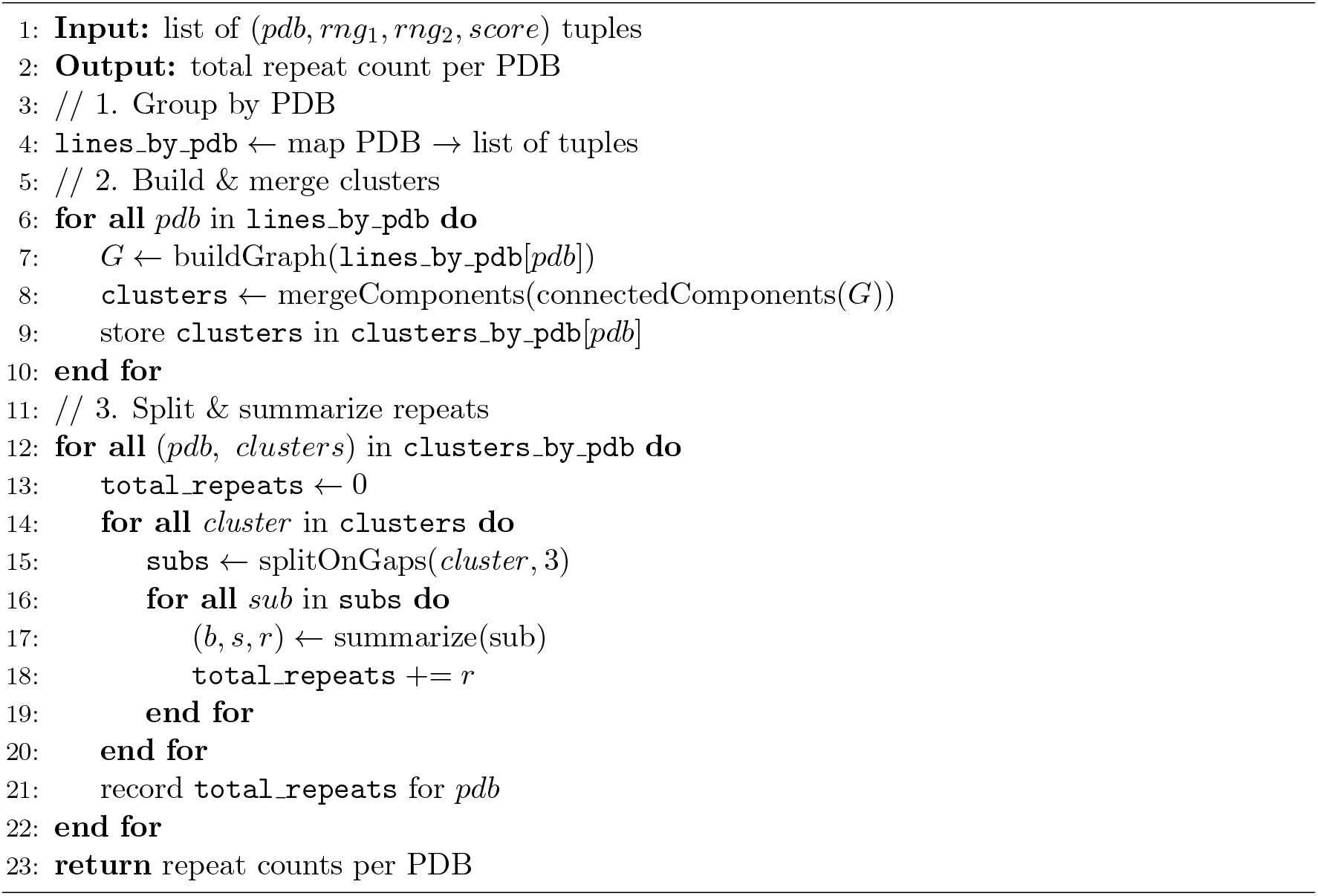

## Code Availability

The source code for the SSE-based structural analysis pipeline is publicly available on GitHub at https://github.com/rl647/SSE_search and data is available at https://github.com/rl647/SSE_search/tree/main/data.

