## Supplementary Information for "Detection of protein symmetry and structural rearrangements using secondary structure elements"

### Contents

- Supplementary Figures

### S1 Supplementary Figures

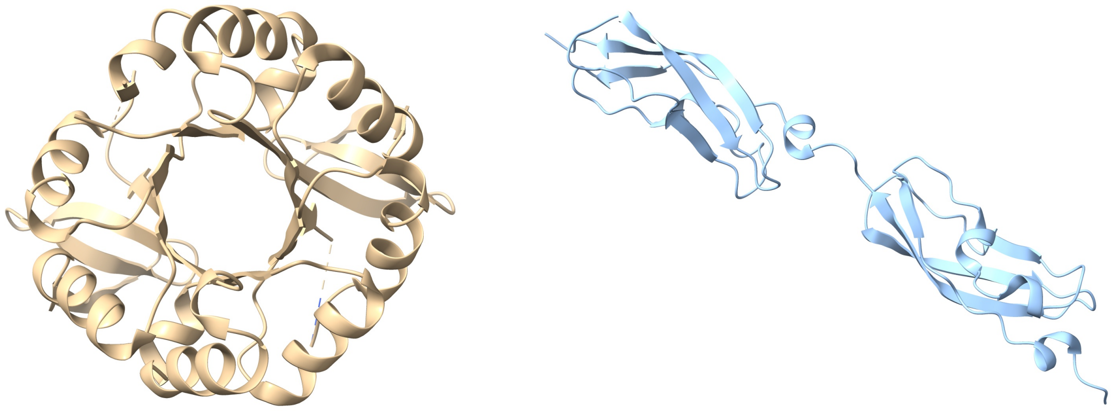

(a) (b)

Figure S1: **Symmetric Proteins Identified by SSEs and CE-Symm.** (a) A protein uniquely identified as symmetric by the SSE-based method, displaying a TM-score of 0.914. (b) A protein uniquely identified by CE-Symm, with a TM-score of 0.960. These examples demonstrate that while both methods capture a comparable number of symmetric proteins, each uniquely detects distinct local symmetry features.

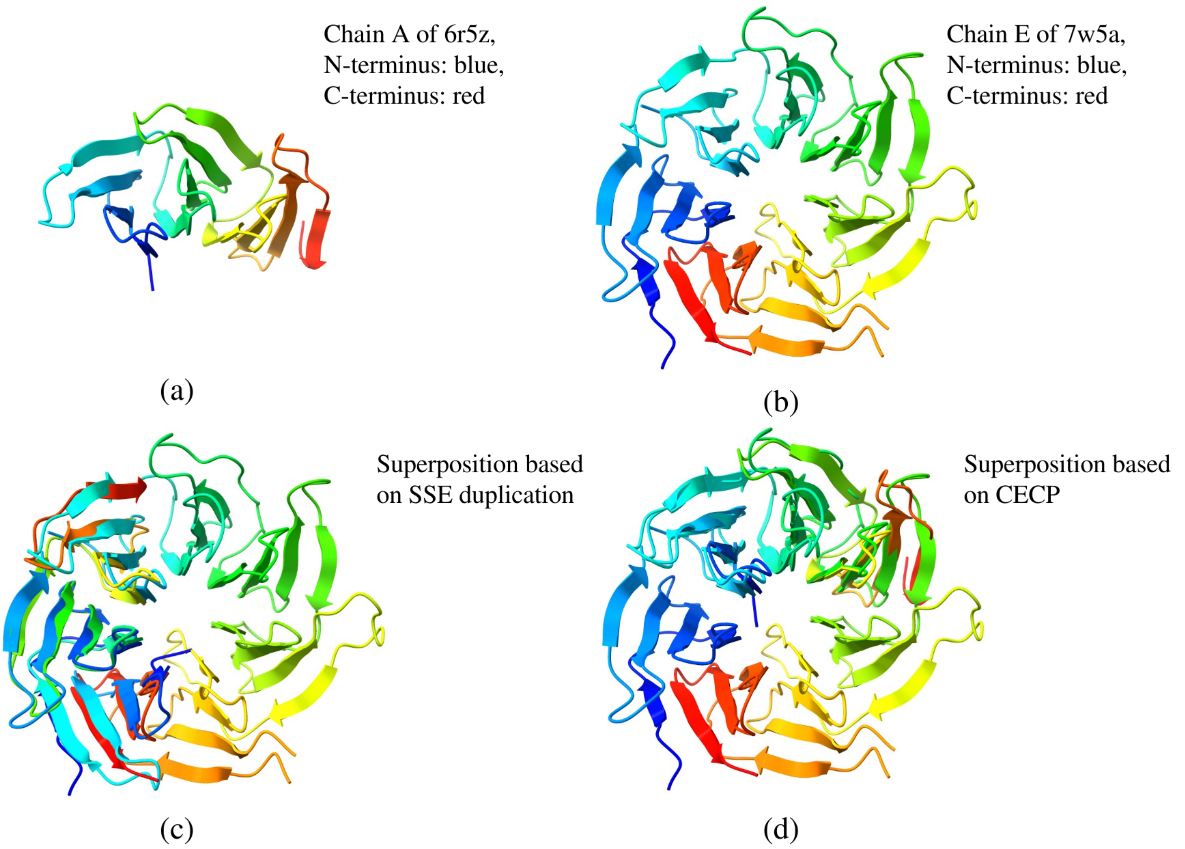


Figure S2: **Example of a circular permutation detected by SSE duplication but not by CECP, with similar TM-scores.** (a) Chain A of PDB 6r5z, colored from blue at the C-terminus to red at the N-terminus; (b) Chain E of PDB 7w5a, colored from blue at the C-terminus to red at the N-terminus; (c) Superposition using the SSE-duplication–derived boundary; (d) Superposition using the CECP-derived boundary. Both them show very well matched which means symmetry is a special type of Circular Permutations.

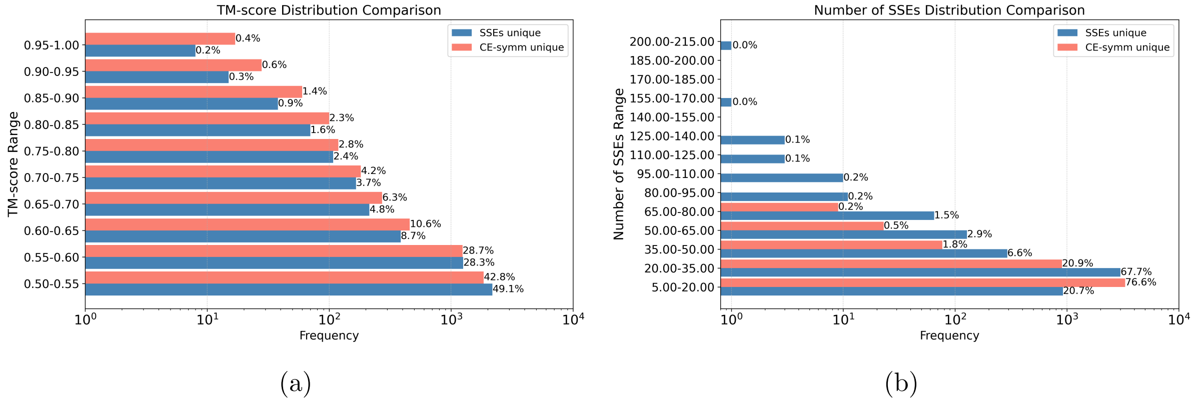


Figure S3: **Comparison of Unique Predictions by SSEs and CE-Symm.** (a) TM-score distribution of unique predictions from SSEs and CE-Symm. Both methods show a similar distribution, with CE-Symm identifying slightly more symmetric proteins when the symmetric units are highly similar. (b) Distribution of the number of SSEs in unique predictions. CE-Symm captures more symmetry in smaller proteins, while SSEs are more effective in identifying symmetry in larger proteins. These results suggest that CE-Symm and SSEs capture complementary aspects of structural symmetry.

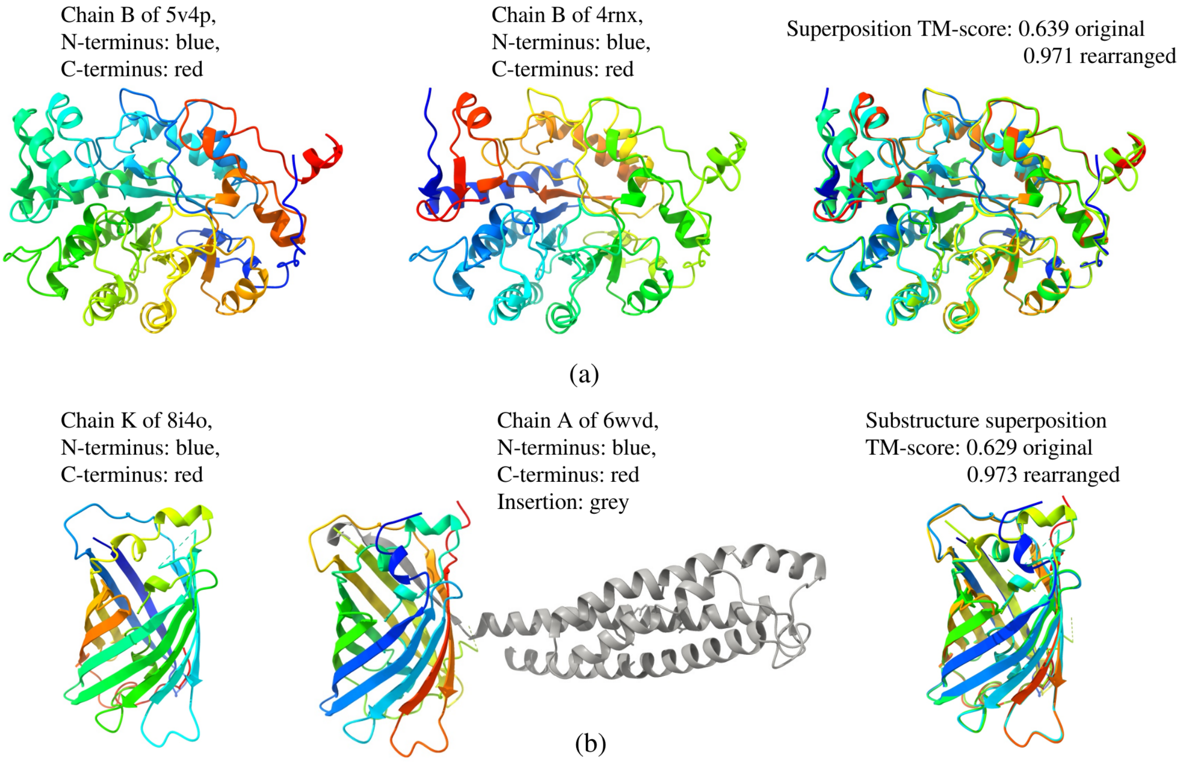


Figure S4: **Structural Clustering of Circular Permutants and Indel Mutants.** (a) Example of circular permutation. (b) Example of circular permutation with indel mutation. These examples demonstrate the effectiveness of the SSE-based approach in detecting structural similarity in circular permutations and the impact of indel mutations on clustering.

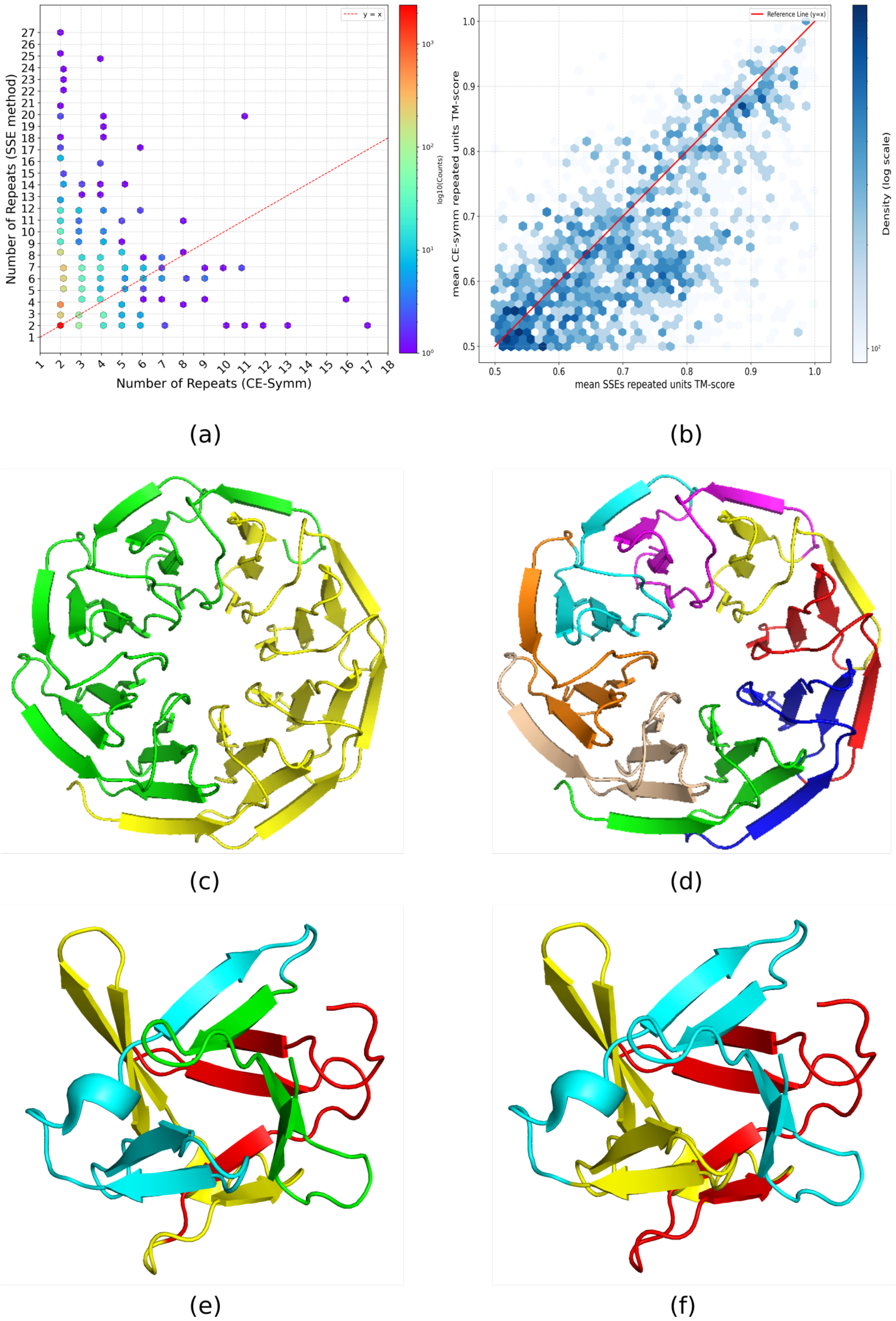


Figure S5: **Comparison of repeated units identified by SSE-based and CE-Symm methods.** (a) Comparison of the number of repeats identified by SSEs and CE-Symm. (b) Comparison of average TM-scores of symmetric units from both methods. (c, d) Symmetric units in chain A of 6tjg: CE-Symm identifies 8-fold whilst SSEs identifies 2-fold. (e, f) Symmetric units in chain A of 3p6j: CE-Symm identifies 3-fold whilst SSEs identifies 4-fold. Overall, the two methods generally agree, though CE-Symm tends to find more symmetric units.
